# Inference and visualization of DNA damage patterns using a grade of membership model

**DOI:** 10.1101/327684

**Authors:** Hussein Al-Asadi, Kushal K Dey, John Novembre, Matthew Stephens

**Affiliations:** Committee on Evolutionary Biology, University of Chicago,Chicago, 60637, United States; Department of Statistics, University of Chicago, Chicago, 60637, United States; Department of Human Genetics, University of Chicago, Chicago,60637, United States

## Abstract

Quality control plays a major role in the analysis of ancient DNA (aDNA). One key step in this quality control is assessment of DNA damage: aDNA contains unique signatures of DNA damage that distinguish it from modern DNA, and so analyses of damage patterns can help confirm that DNA sequences obtained are from endogenous aDNA rather than from modern contamination. Predominant signatures of DNA damage include a high frequency of cytosine to thymine substitutions (C-to-T) at the ends of fragments, and elevated rates of purines (A & G) before the 5’ strand-breaks. Existing QC procedures help assess damage by simply plotting for each sample, the C-to-T mismatch rate along the read and the composition of bases before the 5’ strand-breaks. Here we present a more flexible and comprehensive model-based approach to infer and visualize damage patterns in aDNA, implemented in an R package `aRchaic`. This approach is based on a “grade of membership” model (also known as “admixture” or “topic” model) in which each sample has an estimated grade of membership in each of *K* damage profiles that are estimated from the data. We illustrate `aRchaic` on data from several aDNA studies and modern individuals from 1000 Genomes Project Consortium (2012). Here, `aRchaic` clearly distinguishes modern from ancient samples irrespective of DNA extraction, lab and sequencing protocols. Additionally, through an *in-silico* contamination experiment, we show that the `aRchaic` grades of membership reflect relative levels of exogenous modern contamination. Together, the outputs of `aRchaic` provide a concise visual summary of DNA damage patterns, as well as other processes generating mismatches in the data. **Availability**: `aRchaic` is available for download from https://www.github.com/kkdey/aRchaic.

## 1 Introduction

Ancient DNA (aDNA) research has seen rapid growth with the recent advancements in recovery of short DNA fragments, increased throughput, and lower per-base cost in sequencing (Shapiro and Hofreiter, 2014). Both the number and size of aDNA datasets have grown rapidly over the last five years, and several recent studies sequenced hundreds of ancient individuals (Mathieson et al., 2015; Allentoft et al., 2015; Mathieson et al., 2017; Olalde et al., 2017; Lazaridis et al., 2016; Lipson et al., 2017).

This rapid recent growth in aDNA research has provided novel insights into human history. However, working with aDNA remains challenging. For example, ancient samples often contain very little endogenous DNA because in many environments DNA degrades rapidly post-mortem (Sawyer et al., 2012). Furthermore, ancient samples are often contaminated by microbes and exogenous human DNA (Malmström et al., 2007). Both these factors mean that many sequence reads generated by an aDNA study may not actually come from the ancient sample.

Because of these challenges, aDNA researchers pay careful attention to quality control (QC), including checking sequencing reads from each sample for signatures of endogenous aDNA. These signatures include: short fragment length, an enrichment of purines before strand breaks, and a high frequency of cytosine to thymine substitutions (C-to-T) at the ends of fragments (Sawyer et al., 2012; Ginolhac et al., 2011; Jónsson et al., 2013; Briggs et al., 2007; Skoglund et al., 2014b). One common QC procedure is to plot, for each sample, the C-to-T mismatch rate as a function of position from the end of the read, and to look for an elevated rate near the ends of reads as an indication of the presence of endogenous aDNA. Another common procedure is to look for elevated rates of purines (A & G) before the 5’ strand-breaks. Both these procedures are implemented in the software *mapdamage* (Ginolhac et al., 2011; Jónsson et al., 2013), for example.

These commonly-used QC checks, though simple and useful, have several limitations. For example, they produce a plot for each sample, which can be inconvenient to work with and difficult to compare across many samples. This issue becomes increasingly important with the growing size of aDNA datasets. These plots can also be difficult to interpret, in part because they do not contrast observed patterns with expected patterns in modern samples. Finally, these approaches can detect only pre-defined damage signatures, and may fail to capture other key features or artifacts in the data.

Here we introduce methods to help address these problems. These methods start with a Binary Alignment Map (BAM) file, obtained by aligning each read to a reference genome. The BAM file includes information on the mismatches that occur in each read (vs the reference). We characterize each mismatch by several relevant features, including its type (e.g. C-to-G, etc), flanking bases, and distance from the end of the read. We then use these features to cluster the mismatches into groups, which we call *mismatch profiles*. Intuitively, a mismatch profile associated with post-mortem damage is expected to show high levels of C-to-T mismatches at the ends of reads. On the other hand, a mismatch profile that is typical of modern DNA polymorphism will show a different pattern, such as a transition to transversion ratio of 2:1 (Goldman and Yang, 1994). Finally we estimate the relative frequency of each mismatch profile in each sample, which we refer to as the “Grade of membership” (Erosheva, 2006) of that sample in that mismatch profile. These grades of membership should reflect which processes generated mismatches in each sample. For example ancient samples should have a high grade of membership in mismatch profiles characteristic of post-mortem DNA damage. Grade of membership models are widely used to infer structure in admixed populations (Pritchard et al., 2000), document collections (Blei et al., 2003), RNA-seq data Dey et al. (2017a) and somatic mutation data (Shiraishi et al., 2015) for example.

We have implemented methods to fit this model, and visualize the results in a software package, `aRchaic`. For example, the grades of membership for all samples are succinctly displayed in a single STRUCTURE plot (Rosenberg et al., 2002), and each mismatch profile is displayed using simple intuitive plots (Dey et al., 2017b). Together these plots provide a concise visual summary of DNA damage patterns, as well as other processes generating mismatches in the data.

## 2 Methods

For each sample *i* = 1,…, *I* we first obtain a BAM file. From this BAM file we extract information on the mismatches (vs a reference) that occur in reads from the sample. First we filter out low-quality reads (mapping ≤ 30), low-quality mismatches (base quality ≤ 20), and mismatches that occur more than 20bp from the end of a read (since these are unlikely to reflect damage patterns (Briggs et al., 2007)). When a read carries more than one mismatch we treat these as independent (an assumption we verified by checking that the probability of a mismatch conditional on the occurrence of another mismatch on the same read is not significantly different from the marginal probability, p-value = 0.43).

Let *J*_*i*_ denote the total number of remaining mismatches. For each mismatch *j* = 1, …, *J*_*i*_, we first identify the strand in the reference genome that carries a C or T allele; we call this strand the “reference strand” and denote it by *S*_*j*_. Let *x*_*i,j*_ = (*x*_*i,j*,1_, *x*_*i,j*,2_, *x*_*i,j*,3_, *x*_*i,j*,4_, *x*_*i,j*,5_) denote the following features of the mismatch (see Supplementary Figure S1 for illustration):

1. *x*_*i,j*,1_ ∈ {T-to-A, T-to-C, T-to-G, C-to-A, C-to-G, C-to-T} denotes the mismatch (with respect to the reference strand *S*_*j*_).
2. *x*_*i,j*,2_ ∈ {A, C, G, T} denotes the nucleotide immediately 5’ to the mismatch on *S*_*j*_.
3. *x*_*i,j*,3_ ∈ {A, C, G, T} denotes the nucleotide immediately 3’ to the mismatch on *S*_*j*_.
4. *x*_*i,j*,4_ ∈ {0, …20} denotes the distance (in base-pairs) from the mismatch to the nearest end of the read.
5. *x*_*i,j*,5_ ∈ {A, C, G, T} denotes the nucleotide that occurs one base upstream from the 5’ end of the reference strand that is closest to the location of mismatch. Since mismatches resulting from damage primarily occur at 5’ ends of the reads, this features captures the enrichment of purines immediately one base 5’ upstream of the strand-break for samples with sufficient DNA damage (Sawyer et al., 2012; Briggs et al., 2007).

These features are designed to reflect the major modes of DNA damage (Briggs et al., 2007; Sawyer et al., 2012; Prüfer et al., 2014; Sawyer et al., 2012)). For each feature *l* ∈ {1,…, 5} we let *M*_*l*_ denote the number of possible values of *x*_*i,j,l*_, and for notational convenience we relabel the possible outcomes such that *x*_*i,j,l*_ takes on integer values in {1,…, *M*_*l*_}. For example we represent *x*_*i,j*,1_ = T-to-A by *x*_*i,j*,1_ = 1.

Our model assumes that each mismatch in each individual arose from one of *K* mismatch profiles (“clusters”). We introduce latent variables *z*_*i,j*_ ∈ {1,…, *K*} to denote the profile that gave rise to mismatch *j* in individual *i*. We assume

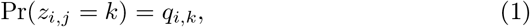

where *q*_*i,k*_ represents the membership proportion of individual *i* in mismatch profile (cluster) *k* ∈ 1,…,*K*.

We further assume that, given *z*_*i,j*_ = *k*,

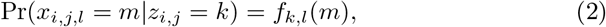

where *m* ∈ {1,…,*M*_*l*_}, and *f*_*k,l*_(*m*) denotes the relative frequency of *m* at feature *l* in cluster *k*. We follow (Shiraishi et al., 2015) in assuming independence among features within each cluster.

Putting this all together, and assuming independence of observations yields the likelihood:

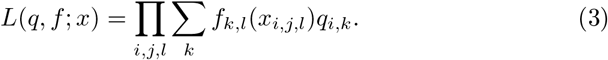

We fit this model, and estimate the individual parameters (*q*) and cluster parameters (*f*) by maximum likelihood using an accelerated EM algorithm. We use the same EM updates as in equations 2-4 in (Shiraishi et al., 2015), and we add first-order quasi-Newton acceleration to improve convergence (Lange, 1995; Alexander et al., 2009; Taddy, 2012).

For each cluster *k*, we visualize the cluster parameters *f*_*k*_ using an *EDLogo* plot (Dey et al., 2017b), see Figure 1. The *EDLogo* plot allows one to visualize both enrichment and depletion of mismatch features scaled against a reference frequency. In our application, the reference frequency was computed by averaging the proportion of the five `aRchaic` features across individuals in the 1000 Genomes Project. This allows us to to compute an enrichment score which effectively compares mismatch profiles in our samples against that of modern individuals from 1000 Genomes Project Consortium (2012). We use a STRUCTURE plot (Rosenberg et al., 2002) to visualize the estimates of *q*_*i,k*_ for each sample.

**Figure 1:**
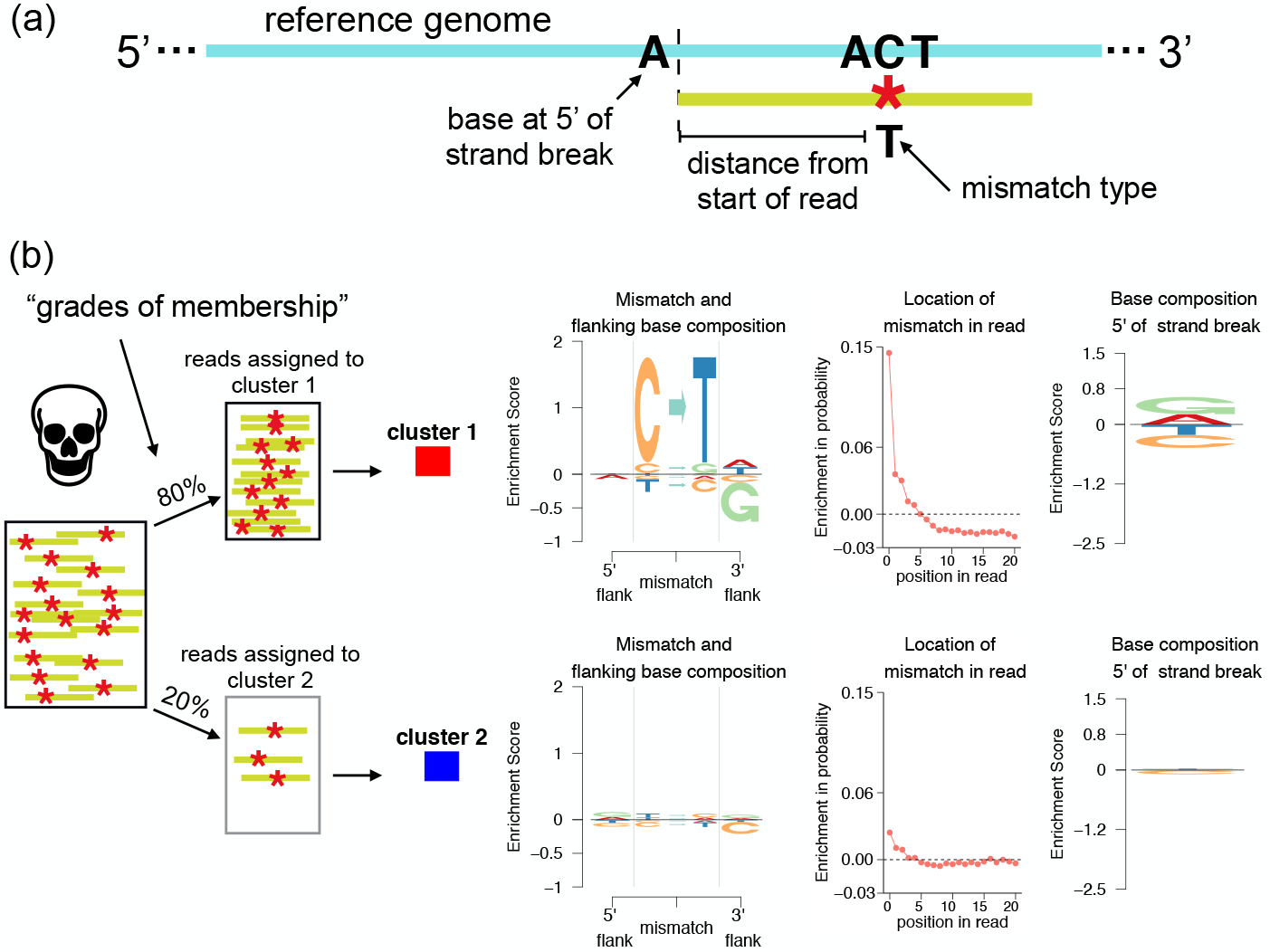
Illustration of the `aRchaic` grades of membership and mismatch profiles. (a) The features of a mismatch modeled by `aRchaic` (b) A depiction of an ancient DNA sample that has 80% of its reads assigned to cluster 1 and 20% of its reads assigned to cluster 2. Each cluster is defined by a *mismatch profile* showing the enrichment of the mismatch type, bases flanking the mismatch, the distance of the mismatch from the nearest end of the read, and the base immediately 5’ to the strand-break. To produce a mismatch profile for a cluster, mismatch features are aggregated across reads assigned to the cluster, and their frequencies are represented by an *EDLogo* plot (Dey et al., 2017b). In the *EDLogo* plot, the frequencies are scaled against a background frequency computed from 1000 Genomes Project Consortium (2012).

## 3 Results

We demonstrate the utility of `aRchaic` using three case-studies.

### 3.1 aRchaic clustering of modern and ancient individuals

We applied `aRchaic` to a combined dataset of 52 ancient samples from four recent studies Skoglund et al. (2014a); Gamba et al. (2014); Lazaridis et al. (2016); Lipson et al. (2017) and 60 modern samples from 1000 Genomes Project Consortium (2012) (n=50) and 10 individuals from HGDP (Cann et al.) individuals sequenced by Meyer et al. (2012). Two of the aDNA studies used partial-UDG treated libraries, which removes most - but not all - of the C-to-T deamination (Rohland et al., 2015).

Figure 2 shows results from `aRchaic` with *K* = 3 (see Supplementary Figure S2 for *K* = 4,5,6). To give a sense of computational requirements, these results took approximately 23 minutes to generate on a single modern compute node. The results clearly highlight differences between modern, ancient (UDG), and ancient (non-UDG) samples. The modern samples show very strong membership in a single cluster (red). As expected, this “modern” cluster shows only modest enrichment in its mismatch type, flanking base composition, and mismatch location, relative to the modern background.

**Figure 2:**
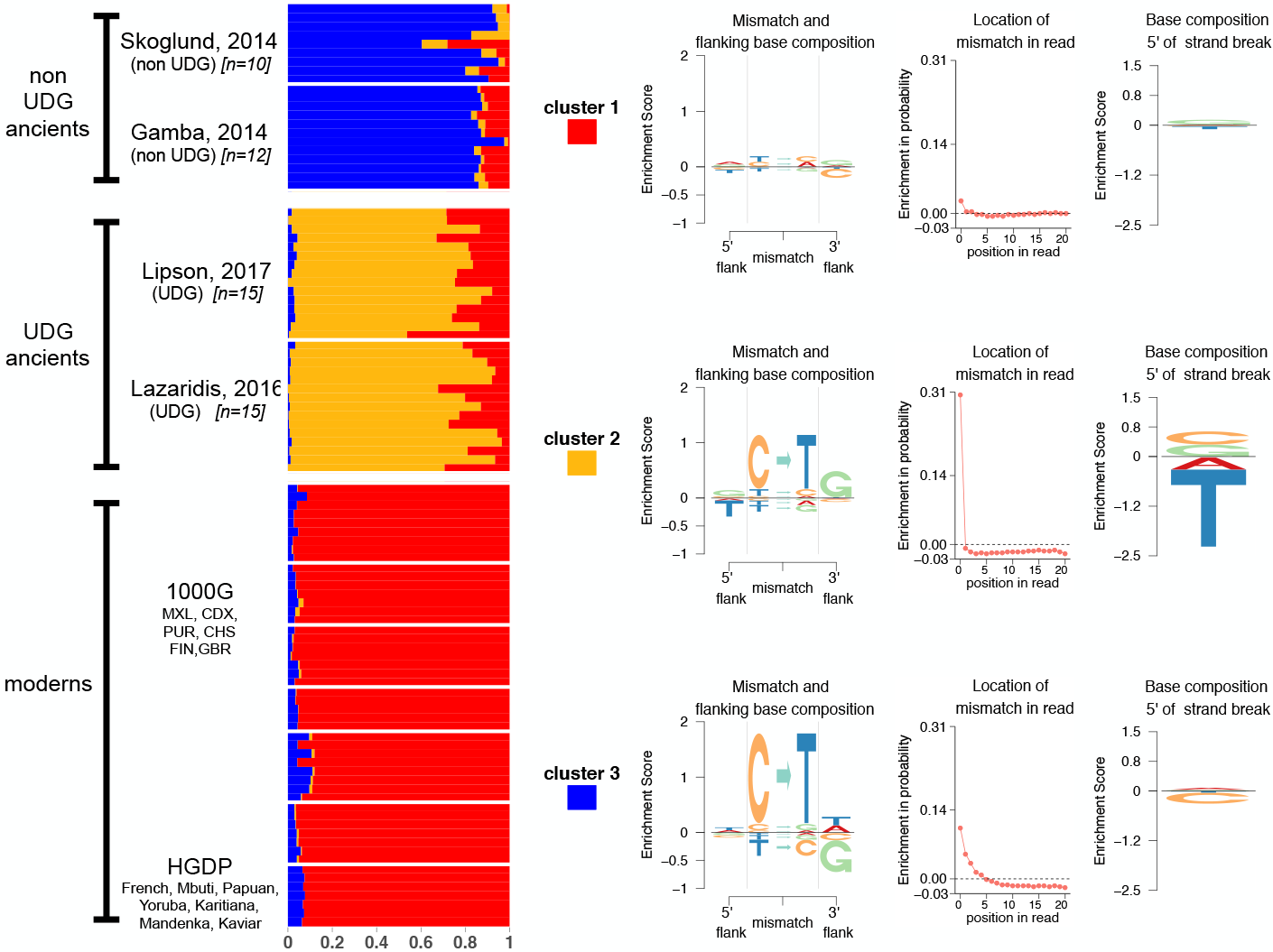
`aRchaic` clearly distinguishes between modern, ancient (UDG), and ancient (non-UDG) samples. `aRchaic` is applied with *K* = 3 to a collection of ancient individuals from four studies Skoglund et al. (2014a); Gamba et al. (2014); Lazaridis et al. (2016); Lipson et al. (2017) along with modern individuals randomly sampled from the 1000 Genomes Project and 10 individuals from the Human Genome Diversity Panel (1000 Genomes Project Consortium, 2012; Cann et al.; Meyer et al., 2012). Modern samples have high membership in the red cluster. The *EDLogo* representation of this cluster does not show strong enrichment against a modern background. The ancient (non-UDG) samples are representative of the blue cluster. The *EDLogo* plot for the blue cluster shows a strong enrichment in C-to-T mismatches at the end of reads, a depletion of guanine in the right flanking base, and a depletion of cytosine at the 5’ strand-break. The ancient (UDG) samples have partial membership both in the red cluster and in the gold cluster. The *EDLogo* plot for the gold cluster is enriched in C-to-T mismatches at the terminal ends of the reads, shows an enrichment of guanine at the right flanking base, and a depletion of thymine one base 5’ upstream of the strand break.

Ancient (non-UDG) samples show high membership in a second cluster (blue). This cluster is characterized by a very strong enrichment of C-to-T mismatches at the ends of the reads, which is a typical sign of DNA damage (Rohland et al., 2015 this enrichment is accompanied by a depletion of guanine just 3’ to the mismatch. We see this depletion in guanine because the blue cluster is driven by mismatches at cytosine sites which seldom precedes a guanine because CpG sites occur less frequently than expected (Shen et al., 1994).

The ancient UDG-treated individuals show high membership in the third (orange) cluster, and partial membership in the first (red) cluster. The membership in the red cluster presumably reflects the fact that the UDG-treatment repairs much of the damage in these samples, making them look more “modern” in their mismatch profiles. The orange cluster is characterized by enrichment of C-to-T mismatches very close to the ends of reads, with a strong enrichment of guanine at the right flanking base. That is, an enrichment of CpG-to-TpG mismatches at the ends of reads. This may be explained by the fact that when a methylated cytosine undergoes deamination it becomes thymine (in contrast to unmethylated cytosines, which deaminate to uracil) and these thymines are not repaired by the UDG-treatment (Duncan and Miller, 1980). Furthermore, we see a depletion of thymine 5’ upstream of the strand-break, which consequently is manifested as a depletion of thymine at the left flanking base.

### 3.2 The effects of contamination on inferred grades of membership

We next sought to examine the effects of exogenous modern contamination on inferred grades of memberships in ancient samples.

We performed an *in-silico* experiment to artificially contaminate ancient samples with modern data from the 1000 Genomes Project (1000 Genomes Project Consortium, 2012). We selected one BAM file from an ancient sample (K01 from (Gamba et al., 2014)), and split its reads into 10 equal subsets. We then contaminated each subset with reads from a distinct modern individual from the 1000 Genomes Project, varying the contamination level from 0% to 100% (Figure 3a). This results in 10 samples (S1-S10) representing 10 contaminated ancient samples with known levels of modern contamination.

**Figure 3:**
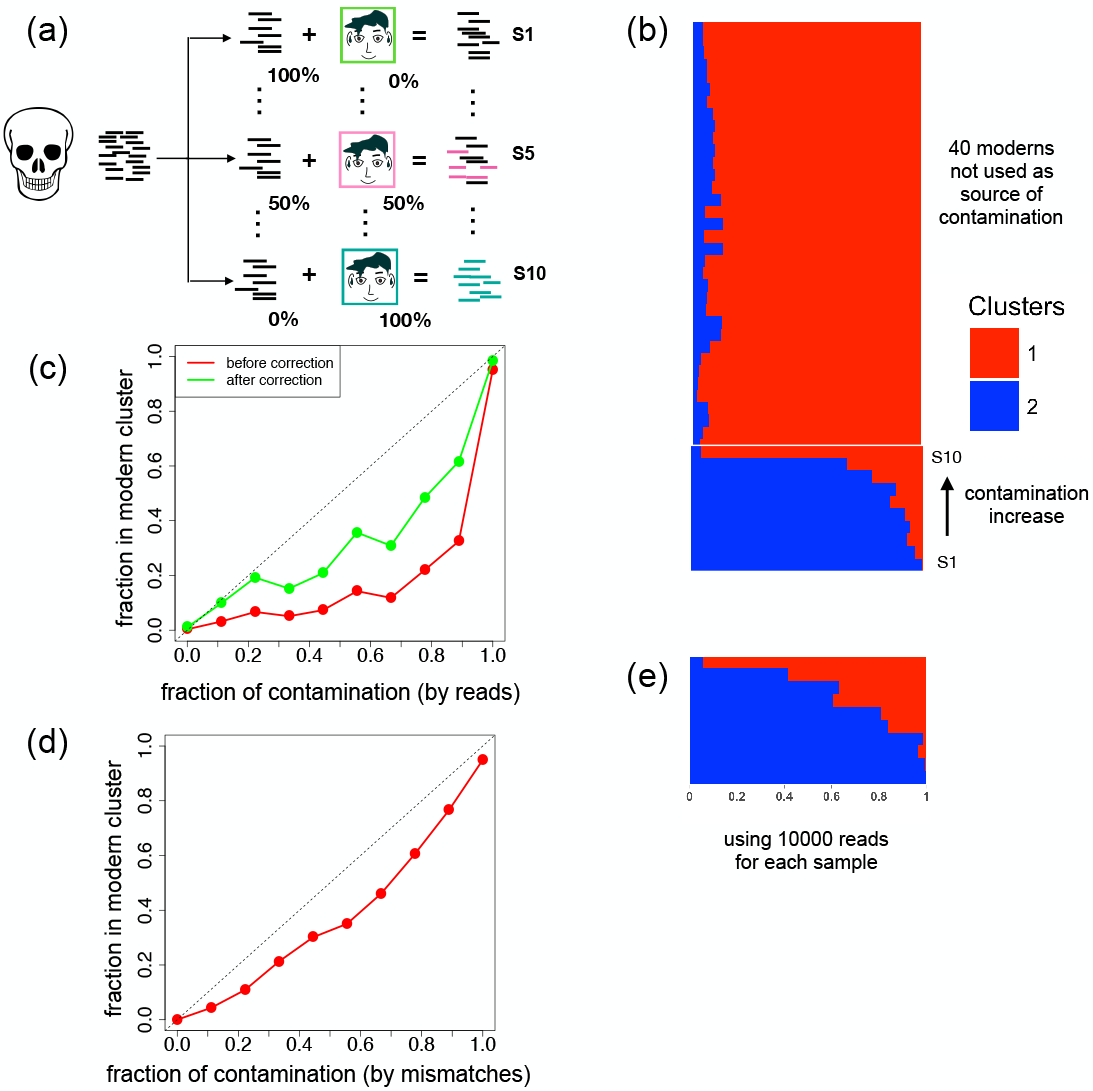
Estimated grades of membership reflect levels of contamination. (a) Reads from one ancient individual (KO1 from Gamba et al. (2014)) were split into 10 equally sized groups. Reads were added from a distinct individual in the 1000 Genomes Project (1000 Genomes Project Consortium, 2012) to each group (S1-S10) at varying levels of percentages (indicating levels of contamination). (b) We applied `aRchaic` with *K* = 2 on a combined dataset comprised of these 10 contaminated groups of reads (S1-S10) along with 40 other modern individuals from 1000 Genomes (c) The grades of membership in cluster 1 (“modern” cluster) were plotted as a function of the percentage of contamination before (red curve) and after (green curve) applying the correction factor discussed in the last paragraph of Section (3.2). (d) We repeated the same experiment as described in panel a, except we discarded all reads with no mismatches or greater than one mismatch. The grades of membership in cluster 1 were plotted as a function of the “mismatch contamination rate” which is defined as the proportion of mismatches that originate from a contaminated read (e) Each group (S1-S10) was further sub-sampled to 10,000 reads, and `aRchaic` was applied with *K* = 2 to the new subsampled groups and the same 40 modern individuals as in panel b.

We applied `aRchaic` with *K* = 2 on the contaminated samples (S1-S10) plus 40 other modern individuals (randomly sampled from the 1000 Genomes Project; Figure 3b). Modern individuals showed high grades of membership in one cluster (red). Fully contaminated individuals nearly showed full membership in the red cluster, while uncontaminated ancient samples showed essentially no membership in this cluster. For samples in between, membership in the red cluster increased approximately monotonically with the level of contamination (Figure 3c). We obtained similar results even with only 10000 randomly-sampled reads for each sample (Figure 3e) implying that these results are robust to low sequencing depth.

We find that the grades of membership in the red cluster (representing moderns) are consistently less than the proportion of contaminated reads (Figure 3c red curve). This pattern can be partly explained by the fact that the vast majority of reads from a DNA library contain no damage. To elaborate, only a fraction of contaminated DNA from a modern source have mismatches, and this fraction will typically be less than for ancient DNA. Thus, the estimated proportion of mismatches arising from a “modern DNA” cluster should generally be expected to be less than the contamination fraction of the library. This implies `aRchaic` will typically provide a lower bound on contamination rates.

In this experiment, if we define the “mismatch contamination rate” as proportion of mismatches that originate from a contaminating read, we see that the `aRchaic` grades of membership are approximately linearly proportional to the mismatch contamination rate, and only slightly underestimate the true proportion of contaminated reads (Figure 3d). This encourages the use of a possible correction factor to infer a true read-level contamination rate. The fraction of a DNA library with mismatches can be measured (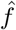, e.g. 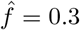), and the fraction of modern DNA reads with mismatches to reference (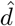, e.g. 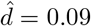) can be approximated using modern samples. With those values a simple moment estimator of the read level contamination rate would be: 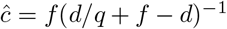, where *q* is the fraction of membership in the “modern” cluster. Applying this estimator to the grade of membership in the the modern cluster in Figure 3c (red curve) provides values that are closer to the true contamination rates (Figure 3c green curve). However, a complication is that high quality aDNA may generate mismatches that cluster with “modern” mismatch profiles even without contamination. This effect is not accounted for by this simple estimator, and is difficult to predict.

### 3.3 aRchaic can identify both DNA damage and technical artifacts

As a final case study, we compiled data from 25 modern and 25 ancient Native Americans from the Northwest Coast of North America (Lindo et al., 2016). This dataset offers us a opportunity to apply `aRchaic` on modern and ancient DNA samples collected from the same population and sequenced in the same laboratory. In these data, the first two positions from the 5’ end of each read had been removed by the original authors in an attempt to mitigate effects of DNA damage. Despite this, `aRchaic`, when applied with *K* = 2 to all 50 samples, clearly distinguishes between modern and ancient individuals (Supplementary Figure S3).

When we applied `aRchaic` to just modern samples we were surprised to find two clear clusters (Figure 4a). These clusters turned out to reflect the fact that the modern samples had been processed using two different library preparation kits, Nextera & TruSeq (Lindo et al., 2016). Samples prepared with the TruSeq kit showed nearly full membership in one cluster (beige), and those prepared with the Nextera kit showed nearly full membership in a second cluster (pink). These clusters show only small differences in mismatch patterns, but the pink cluster is characterized by a strong excess of mismatches at position 12, apparently an artifact introduced by the Nextera preparation (Supplementary Figure S5).

**Figure 4:**
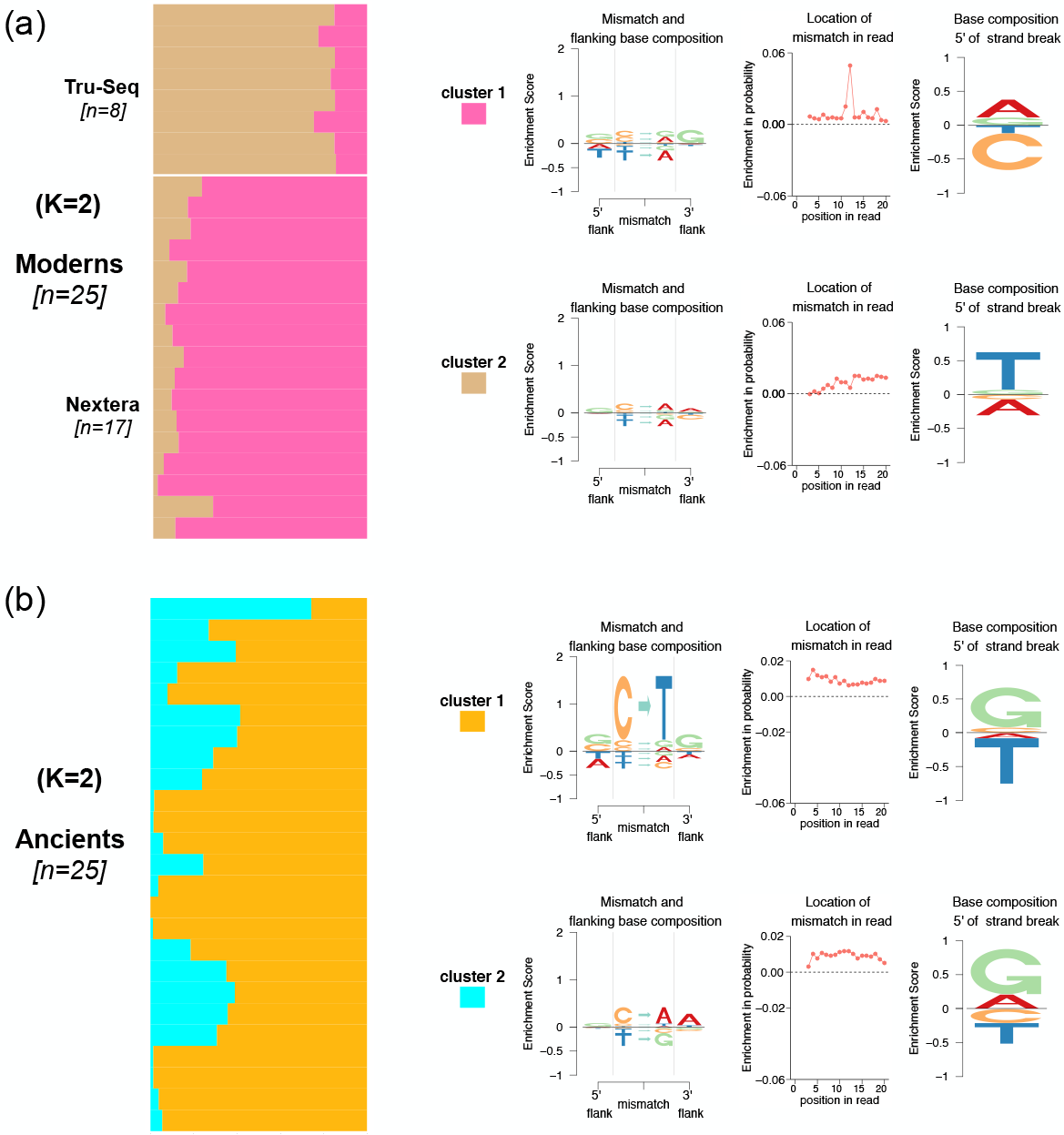
DNA damage and library preparation techniques drive grades of membership. (a) We applied `aRchaic` with *K* = 2 to 25 modern samples from Lindo et al. (2016). The samples prepared with the TruSeq kit show nearly full membership in the pink cluster. Samples prepared with the Nextera kit show partial membership in the pink cluster and the tan cluster. The tan cluster shows a blip at the 12th position from the end of the read (b) We applied `aRchaic` with *K* = 2 to 25 ancient samples from Lindo et al. (2016). The two clusters show an enrichment of C-to-T mismatches at the ends of reads and an enrichment of purines at the 5’ strand-break.

We also applied `aRchaic` with *K* = 2 to just the ancient samples (see Figure 4b). Unlike the moderns, this yielded a continuous gradient of memberships in the two clusters (cyan and gold). One cluster (gold) is dominated by the strong enrichment of C-to-T mismatches relative to the modern background that is typical of ancient samples. The other cluster (cyan) is enriched primarily in C-to-A mismatches, possibly representing another type of damage, or other artifacts, in the ancient DNA. Interestingly, the individual with highest membership in this cyan cluster was much older than all the others (≈ 6000 years BP; all other samples are ≈ 2000 years BP).

## 4 Discussion

We developed a method (`aRchaic`) for clustering and visualization of samples based on DNA mismatch patterns. Our method is based on a Grade-of-Membership (GoM) model, which generalizes the concept of clustering to allow samples to have membership in multiple clusters. We provide a visual representation of the grades of membership using a “Structure plot” (Rosenberg et al., 2002) and visualization of the mismatch profiles (or clusters) with an *EDLogo* plot (Dey et al., 2017b).

In GoM models, the choice of the number of clusters *K* (or mismatch profiles) is a contentious issue. In our analyses we selected values of *K* that highlight interpretable structure in the data. We emphasize that there will typically be no single “true” *K*, and that examining results with different *K* can often provide additional insights (Dey et al., 2017a). For example, Figure 2 shows results for *K* = 3, but higher values of *K* reveal additional structure within each ancient subgroup (Supplementary Figure S2). Similarly, Supplementary Figure S3 shows results for *K* = 2 where the model fails to distinguish between Nextera and TruSeq modern samples, but analysis with *K* = 6 does pick up this difference (Supplementary Figure S4). More generally, `aRchaic` is useful for detecting batch effects as Figure 4 and Supplementary Figure S4 suggest.

A key challenge in analyzing ancient DNA is that data are often contaminated with exogenous modern DNA. Several approaches have been suggested to estimate the amount of contamination. One approach is to compute the rate of polymorphism across the X chromosome in males (Korneliussen et al., 2014; Rasmussen et al., 2011), where the presence of polymorphism would suggest contamination because males have only one X chromosome. Another approach is to quantify the contribution of a panel of modern mitochondrial haplotypes to the ancient DNA (Renaud et al., 2015; Fu et al., 2014). Neither of these approaches leverages autosomal DNA. Although `aRchaic` does not provide explicit estimates of contamination levels, in some settings (e.g. Figure 3) the inferred grades of membership can reflect relative levels of contamination even with low sequencing depth, and may be a useful complement to these other methods. Some caution is necessary though. Under conditions where an ancient sample has undergone a noticeable amount of DNA damage, we underestimate the proportion of contamination (Figure 3). On the other hand, when ancient mismatches are indistinguishable from modern, the proportion in modern cluster of a sample might be greater than the proportion of contamination (as in the case of an ancient sample with little to no damage). We encourage users to keep these caveats in mind, and suggest that in practice `aRchaic` will be most useful for flagging potentially contaminated samples as those that have an above average clustering with modern samples.

Here we have chosen to model features at the level of mismatches, which have been shown to be informative of DNA damage in previous studies (Ginolhac et al., 2011; Jónsson et al., 2013; Briggs et al., 2007). Alternatively, one could formulate a model at the level of reads. For example, the method *PMDtools* computes a score for every read representing the probability that the read is damaged (Skoglund et al., 2014b). This method models mismatches along the read; additionally, one can incorporate indels and fragment length along with mismatches. One reason we chose not to model these extra features was to reduce the feature and computational complexity of our method. Furthermore, these extra features may not actually be driven by DNA damage. For example, we explored fragment length profiles in several aDNA data-sets and found their distributions to be primarily driven by lab-specific effects rather than DNA damage. Another limitation of fragment length is that it can be used only in studies using paired-end sequencing.

Methods described here are available in an open-source R software package at www.github.com/kkdey/aRchaic.

## Acknowledgements

We thank John Lindo for insightful discussions on ancient DNA and DNA damage in particular. We thank the Metlakatla and Lax Kw’alaams First Nations for access to data. The communities are located in the Prince Rupert Harbour region of British Columbia. We thank Anna Gosling, Pontus Skoglund, David Witonsky, Choongwon Jeong, Anna Di Rienzo, and members of the Stephens and Novembre lab for helpful discussions.

## Funding

This work was supported by NIH funding (U01CA198933) to H.A., M.S., and J.N, and the NSF GRFP for H.A.

## Supplementary Figures

**Figure S1:**
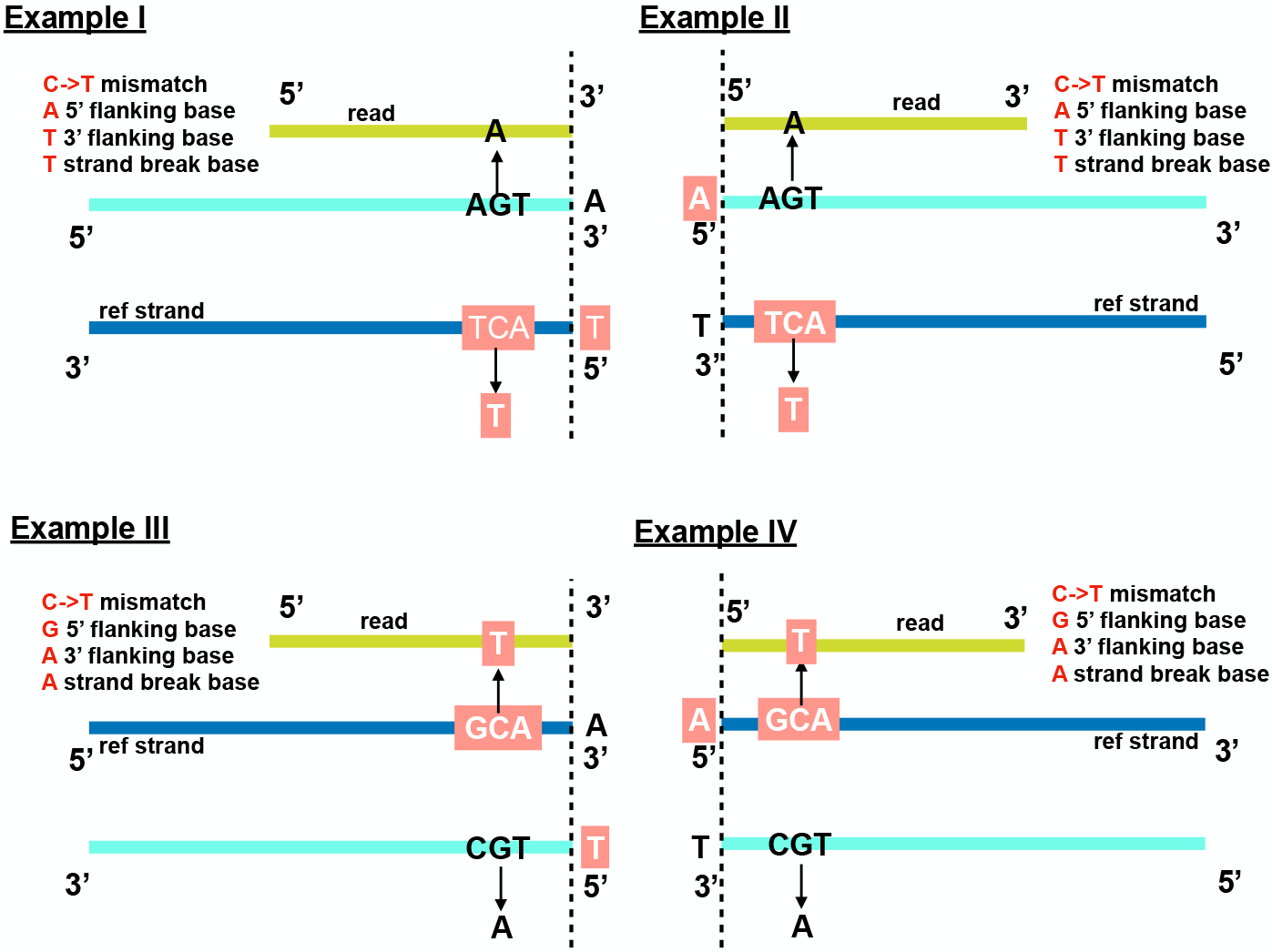
Four examples of how the reference strand for a mismatch is designated. The dark yellow line denotes the mapped read, and the blue and teal line represent the reference genome at two different strand orientations. The reference strand is always designated as the strand that contains the C mismatch or T mismatch (latter not shown).

**Figure S2:**
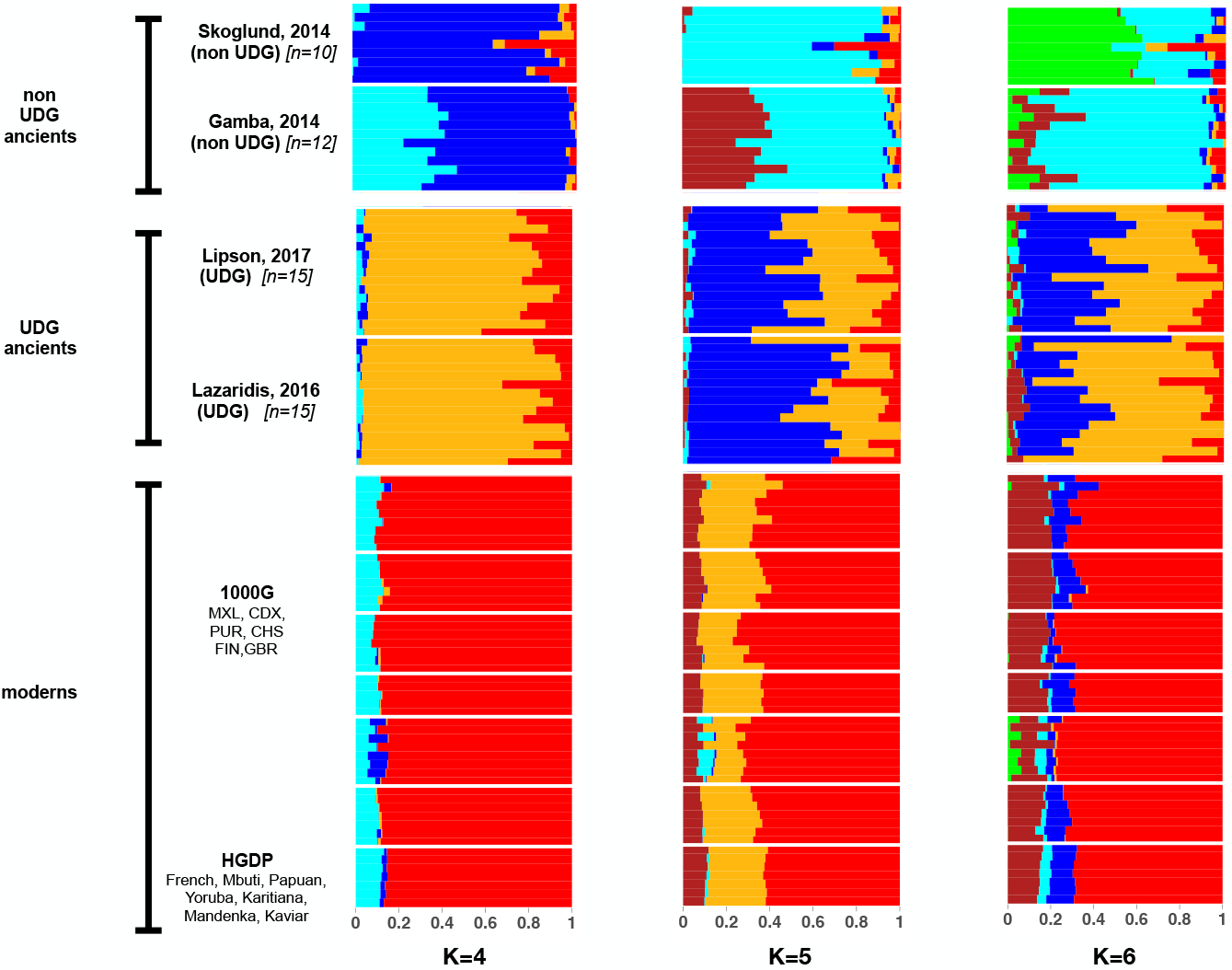
`aRchaic` grades of membership for the example in Fig 1 corresponding to 3 different values of *K* (*K* = 4,5,6). Higher values of *K* distinguish among the ancient studies, reflecting lab and study specific biases.

**Figure S3:**
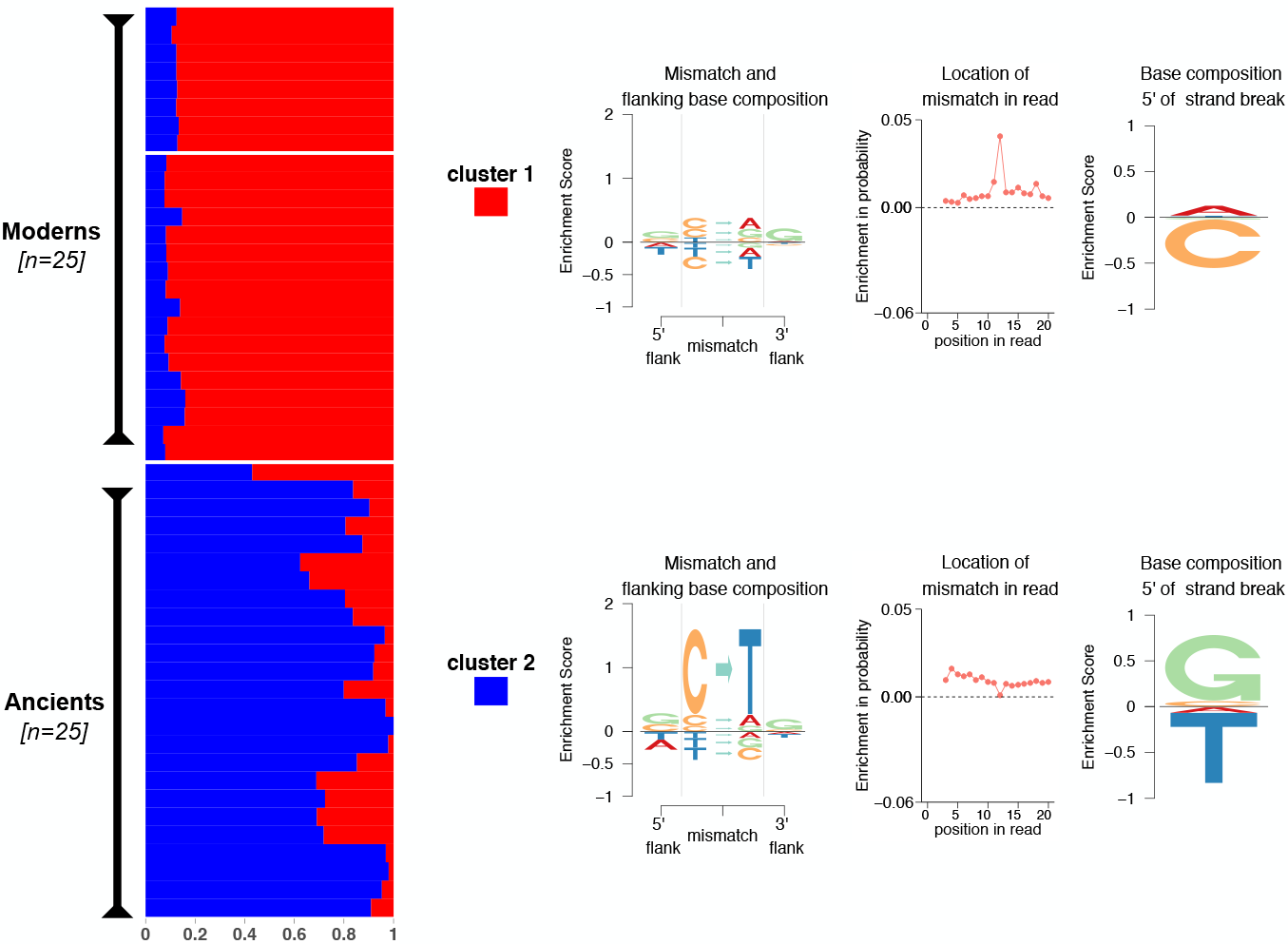
`aRchaic` plot for *K* = 2 on the combined data of 25 moderns and 25 ancients from Lindo et al. (2016). `aRchaic` clearly distinguishes the moderns from the ancients. The ancients are primarily presented by the blue cluster. This cluster shows an enrichment of C-to-T mismatches and depletion of T-to-C mismatches with respect to modern background, as well as enrichment of G and depletion of T at the 5’ strand break. The red cluster shows a blip at 12th position from the end of the read, the explanation for which is provided in Figure S5.

**Figure S4:**
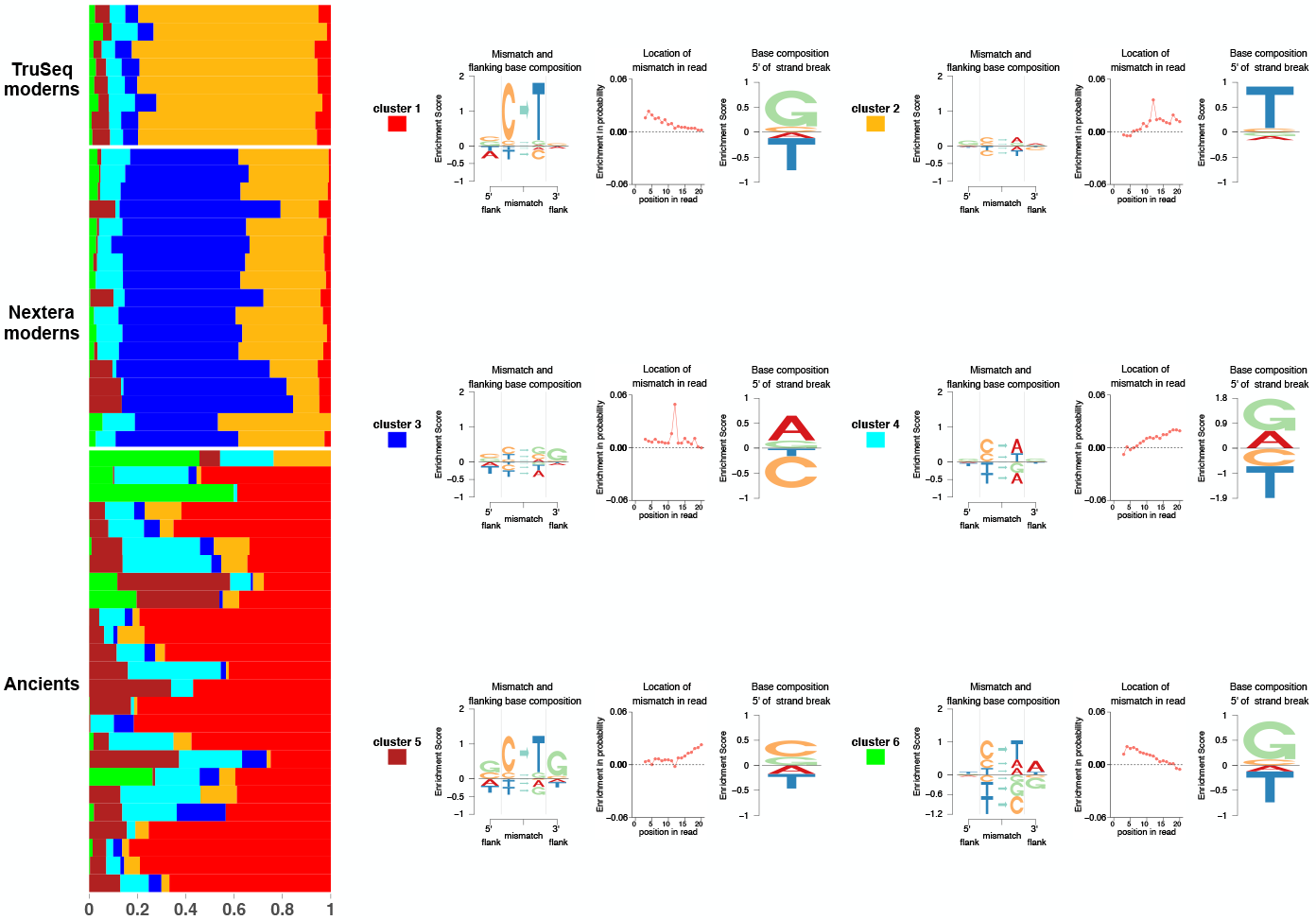
We apply `aRchaic` with *K* = 6 on the data from Fig 4. In addition to separating out the ancients from the moderns, `aRchaic` now distinguishes between moderns individuals based on library kit (Nextera vs Tru-seq.)

**Figure S5:**
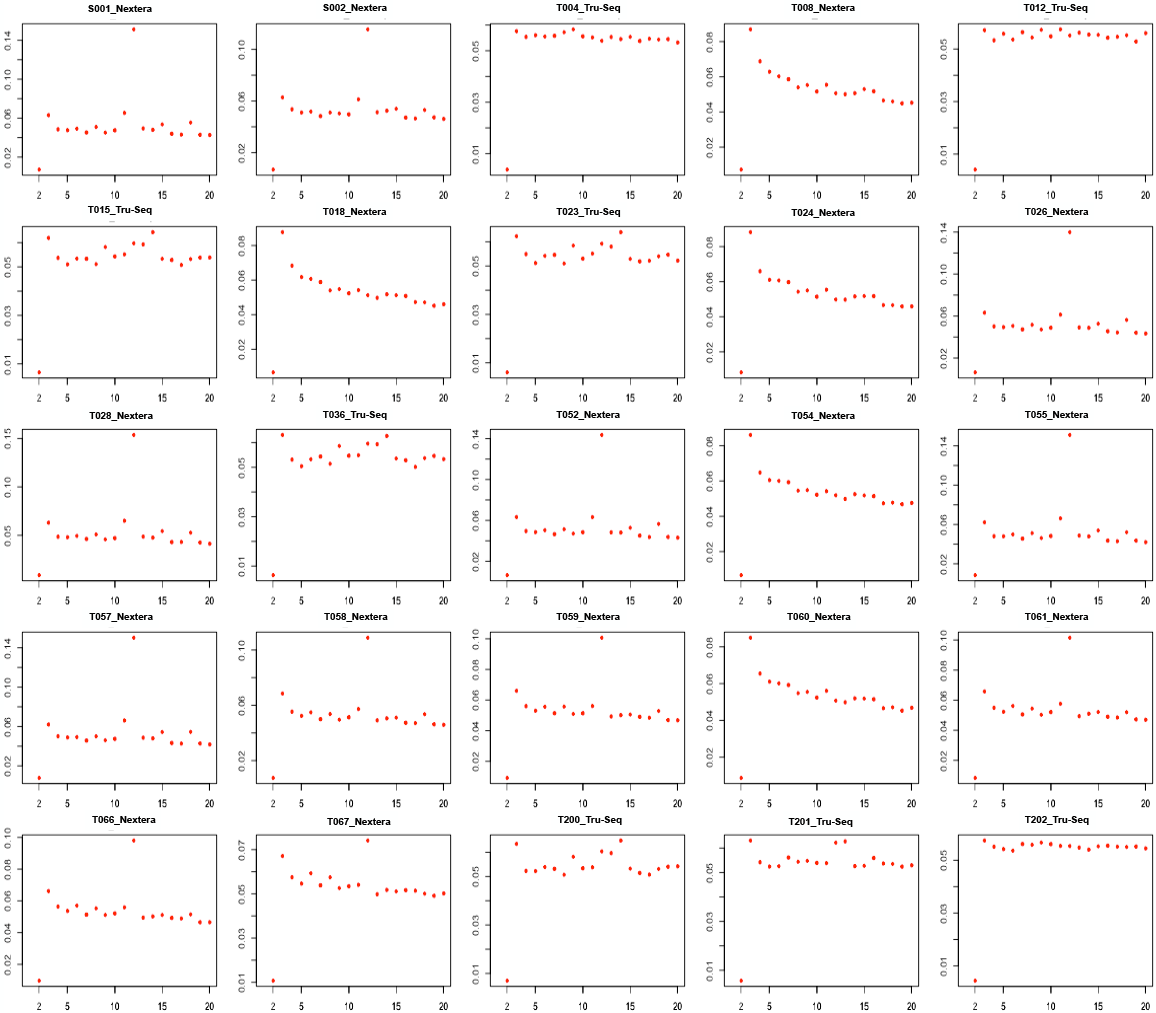
The frequency of all mismatch types plotted against the position of the read (from the 5’ end) for each of the 25 moderns samples in Lindo et al. (2016). Each sample was prepared by one of two library kits: Nextera and TruSeq. Most of the samples prepared with the Nextera kit show a spike in frequency at the 12th position from the 5’ end of the read

## References

1000 Genomes Project Consortium. An Integrated Map of Genetic Variation from 1,092 Human Genomes. Nature, 491(7422):56, 2012.

David H Alexander, John Novembre, and Kenneth Lange. Fast Model-Based Estimation of Ancestry in Unrelated Individuals. Genome research, 19(9): 1655–1664, 2009.

Morten E Allentoft, Martin Sikora, Karl-Göran Sjögren, Simon Rasmussen, Morten Rasmussen, Jesper Stenderup, Peter B Damgaard, Hannes Schroeder, Torbjörn Ahlström, Lasse Vinner, et al. Population Genomics of Bronze Age Eurasia. Nature, 522(7555):167–172, 2015.

David M Blei, Andrew Y Ng, and Michael I Jordan. Latent Dirichlet Allocation. Journal of machine Learning research, 3(Jan):993–1022, 2003.

Adrian W Briggs, Udo Stenzel, Philip LF Johnson, Richard E Green, Janet Kelso, Kay Prüfer, Matthias Meyer, Johannes Krause, Michael T Ronan, Michael Lachmann, et al. Patterns of Damage in Genomic DNA sequences from a Neandertal. Proceedings of the National Academy of Sciences, 104(37): 14616–14621, 2007.

Howard Cann, Claudia De Toma, Lucien Cazes, Marie-Fernande Legrand, Valerie Morel, Laurence Piouffre, Julia Bodmer, Walter F Bodmer, Batsheva Bonne-Tamir, and Anne Cambon-Thomsen. et al A Human Genome Diversity Cell Line Panel.

Kushal K Dey, Chiaowen Joyce Hsiao, and Matthew Stephens. Visualizing the Structure of RNA-seq Expression Data using Grade of Membership Models. PLoS genetics, 13(3):e1006599, 2017a.

Kushal K Dey, Dongyue Xie, and Matthew Stephens. A New Sequence Logo Plot to Highlight Enrichment and Depletion. bioRxiv, 2017b. doi: 10.1101/226597. URL https://www.biorxiv.org/content/early/2017/11/29/226597.

Bruce K Duncan and Jeffrey H Miller. Mutagenic Deamination of Cytosine Residues in DNA. Nature, 287(5782):560, 1980.

Elena A Erosheva. Latent Class Representation of the Grade of Membership Model. Seatle: University of Washington, 2006.

Qiaomei Fu, Heng Li, Priya Moorjani, Flora Jay, Sergey M Slepchenko, Aleksei A Bondarev, Philip LF Johnson, Ayinuer Aximu-Petri, Kay Prüfer, Cesare de Filippo, et al. Genome Sequence of a 45,000-year-old Modern Muman from Western Siberia. Nature, 514(7523):445–449, 2014.

Cristina Gamba, Eppie R Jones, Matthew D Teasdale, Russell L McLaughlin, Gloria Gonzalez-Fortes, Valeria Mattiangeli, László Domboróczki, Ivett Kővári, Ildikó Pap, Alexandra Anders, et al. Genome Flux and Stasis in a Five Millennium Transect of European Prehistory. Nature communications, 5:52–57, 2014.

Aurelien Ginolhac, Morten Rasmussen, M Thomas P Gilbert, Eske Willerslev, and Ludovic Orlando. mapDamage: Testing for Damage Patterns in Ancient DNA Sequences. Bioinformatics, 27(15):2153–2155, 2011.

Nick Goldman and Ziheng Yang. A Codon-based Model of Nucleotide Substitution for Protein-coding DNA Sequences. Molecular biology and evolution, 11(5):725–736, 1994.

Hákon Jónsson, Aurélien Ginolhac, Mikkel Schubert, Philip LF Johnson, and Ludovic Orlando. mapDamage2.0: Fast Approximate Bayesian Estimates of Ancient DNA Damage Parameters. Bioinformatics, 29(13):1682–1684, 2013.

Thorfinn Sand Korneliussen, Anders Albrechtsen, and Rasmus Nielsen. ANGSD: Analysis of Next Generation Sequencing Data. BMC bioinformatics, 15(1): 356, 2014.

Kenneth Lange. A Quasi-Newton Acceleration of the EM Algorithm. Statistica sinica, pages 1–18, 1995.

Iosif Lazaridis, Dani Nadel, Gary Rollefson, Deborah C Merrett, Nadin Rohland, Swapan Mallick, Daniel Fernandes, Mario Novak, Beatriz Gamarra, Kendra Sirak, et al. Genomic Insights into the Origin of Farming in the Ancient Near East. Nature, 536(7617):419, 2016.

John Lindo, Emilia Huerta-Sánchez, Shigeki Nakagome, Morten Rasmussen, Barbara Petzelt, Joycelynn Mitchell, Jerome S Cybulski, Eske Willerslev, Michael DeGiorgio, and Ripan S Malhi. A Time Transect of Exomes from a Native American Population before and after European Contact. Nature communications, 7:13175, 2016.

Mark Lipson, Anna Szécsényi-Nagy, Swapan Mallick, Annamária Pósa, Balázs Stégmár, Victoria Keerl, Nadin Rohland, Kristin Stewardson, Matthew Ferry, Megan Michel, et al. Parallel Palaeogenomic Transects Reveal Complex Genetic History of Early European Farmers. Nature, 551(7680):368, 2017.

Helena Malmström, Emma M Svensson, M Thomas P Gilbert, Eske Willerslev, Anders Götherström, and Gunilla Holmlund. More on Contamination: the Use of Asymmetric Molecular Behavior to Identify Authentic Ancient Human DNA. Molecular biology and evolution, 24(4):998–1004, 2007.

Iain Mathieson, Iosif Lazaridis, Nadin Rohland, Swapan Mallick, Nick Patterson, Songül Alpaslan Roodenberg, Eadaoin Harney, Kristin Stewardson, Daniel Fernandes, Mario Novak, et al. Genome-wide Patterns of Selection in 230 Ancient Eurasians. Nature, 528(7583):499–503, 2015.

Iain Mathieson et al. The Genomic History Of Southeastern Europe. bioRxiv, 2017. doi: 10.1101/135616. URL https://www.biorxiv.org/content/early/2017/09/19/135616.

Matthias Meyer, Martin Kircher, Marie-Theres Gansauge, Heng Li, Fernando Racimo, Swapan Mallick, Joshua G Schraiber, Flora Jay, Kay Prüfer, Cesare De Filippo, et al. A High-coverage Genome Sequence from an Archaic Denisovan Individual. Science, 338(6104):222–226, 2012.

Iñigo Olalde et al. The Beaker Phenomenon And The Genomic Transformation Of Northwest Europe. bioRxiv, 2017. doi: 10.1101/135962. URL https://www.biorxiv.org/content/early/2017/05/09/135962.

Jonathan K Pritchard, Matthew Stephens, and Peter Donnelly. Inference of Population Structure using Multilocus Genotype Data. Genetics, 155(2):945–959, 2000.

Kay Prüfer, Fernando Racimo, Nick Patterson, Flora Jay, Sriram Sankararaman, Susanna Sawyer, Anja Heinze, Gabriel Renaud, Peter H Sudmant, Cesare De Filippo, et al. The Complete Genome Sequence of a Neandertal from the Altai Mountains. Nature, 505(7481):43, 2014.

Morten Rasmussen, Xiaosen Guo, Yong Wang, Kirk E Lohmueller, Simon Rasmussen, Anders Albrechtsen, Line Skotte, Stinus Lindgreen, Mait Metspalu, Thibaut Jombart, et al. An Aboriginal Australian Genome Reveals Separate Human Dispersals into Asia. Science, 334(6052):94–98, 2011.

Gabriel Renaud, Viviane Slon, Ana T Duggan, and Janet Kelso. Schmutzi: Estimation of Contamination and Endogenous Mitochondrial Consensus Calling for Ancient DNA. Genome biology, 16(1):224, 2015.

Nadin Rohland, Eadaoin Harney, Swapan Mallick, Susanne Nordenfelt, and David Reich. Partial uracil-DNA-glycosylase Treatment for Screening of Ancient DNA. Phil. Trans. R. Soc. B, 370(1660):20130624, 2015.

Noah A Rosenberg, Jonathan K Pritchard, James L Weber, Howard M Cann, Kenneth K Kidd, Lev A Zhivotovsky, and Marcus W Feldman. Genetic Structure of Human Populations. science, 298(5602):2381–2385, 2002.

Susanna Sawyer, Johannes Krause, Katerina Guschanski, Vincent Savolainen, and Svante Pääbo. Temporal Patterns of Nucleotide Misincorporations and DNA Fragmentation in Ancient DNA. PloS one, 7(3):e34131, 2012.

B Shapiro and Michael Hofreiter. A Paleogenomic Perspective on Evolution and Gene Function: New Insights from Ancient DNA. Science, 343(6169): 1236573, 2014.

Jiang-Cheng Shen, William M Rideout III, and Peter A Jones. The Rate of Hydrolytic Deamination of 5-Methylcytosine in Double-Stranded DNA. Nucleic acids research, 22(6):972–976, 1994.

Yuichi Shiraishi, Georg Tremmel, Satoru Miyano, and Matthew Stephens. A Simple Model-Based Approach to Inferring and Visualizing Cancer Mutation Signatures. PLoS genetics, 11(12):e1005657, 2015.

Pontus Skoglund, Helena Malmström, Ayça Omrak, Maanasa Raghavan, Cristina Valdiosera, Torsten Günther, Per Hall, Kristiina Tambets, Jüri Parik, Karl-Göran Sjögren, et al. Genomic Diversity and Admixture Differs for Stone-Age Scandinavian Foragers and Farmers. Science, 344(6185):747–750, 2014a.

Pontus Skoglund, Bernd H Northoff, Michael V Shunkov, Anatoli P Derevianko, Svante Pääbo, Johannes Krause, and Mattias Jakobsson. Separating Endogenous Ancient DNA from Modern Day Contamination in a Siberian Neandertal. Proceedings of the National Academy of Sciences, 111(6):2229–2234, 2014b.

Matt Taddy. On Estimation and Selection for Topic Models. In International Conference on Artificial Intelligence and Statistics, pages 1184–1193, 2012.

